# Probing Loop-Mediated Isothermal Amplification (LAMP) targeting two gene-fragments of rose rosette virus

**DOI:** 10.1101/2021.08.17.456656

**Authors:** Andrea Salazar, Francisco M. Ochoa-Corona, Jennifer D. Olson, Binoy Babu, Mathews Paret

## Abstract

This study explores the development of Loop-mediated isothermal amplification of DNA (LAMP) for detection of rose rosette emaravirus (RRV), a technique with the potential to be translated to rose nurseries. RRV is a negative-sense single-stranded RNA *Emaravirus* and causal agent of the rose rosette disease (RRD). Transmission of RRV is by *Phyllocoptes fructiphilus*, an eriophyid mite. Although RRV symptoms are characteristics, early visual diagnosis of RRD can be misleading and confusing since it may appear similar to herbicide damage. Two sets of RRV gene sequences composed of twenty-two accessions of RRV-P3 (RNA 3) and another twenty-four from RRV-P4 (RNA 4) were analyzed and two sets of four LAMP primers were designed for broad-range detection of RRV isolates. The direct antigen-capture method for direct trapping of RRV in plastic was used for RNA extraction followed by cDNA synthesis. LAMP reactions were optimized for Bst 2.0 DNA polymerase using the outer RRV-F3/RRV-B3 primers, and internal RRV-FIP/RRV-BIP primers. LAMP reactions were for 1 hour at 64°C (RRV-P3) and 66.5°C (RRV-P4) using either a thermocycler or a portable dry bath. LAMP was also optimized using DNA polymerase GspSSD LD using the same RRV sets of primers. RRV was detected in symptomatic and non-symptomatic RRD tissue from Oklahoma. The limit of detection (LoD) using Bst 2.0 LAMP was 1pg/μL and 1 fg/μL with GspSSD LD quantitative LAMP. The LoD of pre-reaction hydroxy naphthol blue (HNB, 120 μM) for colorimetric (visual) reactions was 10 pg/μL and 0.1 pg/μL using SYBR green I (1:10 dilution) in colorimetric post-reactions. No cross-reactivity was detected in LAMP reaction testing cDNAs of eight commonly co-infecting rose viruses (INSV, ArMV, MSpV, TSWV, ApMV, PNRSV, ToRSV, and TMV), and one virus taxonomically related to RRV (HPWMoV). RNA from healthy rose tissues and non-template controls (water) were included in all LAMP assays.

## Introduction

Roses are affected by a wide variety of fungi, bacteria, viruses, nematodes, and phytoplasmas, which cause growth reduction, defoliation, and plant death [1]. In most cases, these pathogens attack flowers or leave tissue causing economic losses to landscapers, gardeners, public gardens, and nurseries [1]. In the United States, the total of shrub rose wholesale was 28 million [2]. Rose rosette disease (RRD) is one of the most devastating diseases of roses. This disease was reported in 1940 in Manitoba, Canada, and it has since spread to roses in Kansas, Oklahoma, Missouri, and Arkansas [3]. Currently, RRD is endemic in the north-central, south-central, and southeastern regions of the U.S. [4, 5]. In 2012, thousands of roses at the Tulsa municipal rose garden expressed characteristic symptoms of the disease and were eliminated after the damage inflicted by the disease and as a precautionary sanitary practice to maintain the beauty of the garden [6]. Since then, the rose garden never recovered, all roses got lost and the garden lots have been used to assess rose varietal resistance to RRD.

Rose rosette emaravirus (RRV) is a negative-sense single-strand RNA virus in the genus *Emaravirus*, Bunyaviridae [4]. The eriophyid mite *Phyllocoptes fructiphilus* is the known vector that transmits RRV [4, 7]. RRV appeared in populations of wild roses in the U.S., and it extended to multiflora rose (*Rosa multiflora*), which is a susceptible host. This rose species is invasive and was introduced from Asia, mainly from Japan [8]. RRV has seven genomic RNA segments: RNA1 (RNA-dependent RNA polymerase), RNA2 (glycoprotein), RNA3 (nucleocapsid protein), RNA4 (movement protein), and RNA5, RNA6, and RNA7 specific functions are yet to be determined [4, 9, 10]. RRD symptoms are observed from mid-spring to fall; symptom expression depends mainly on the rose cultivar, age of the plant, and growing conditions. The RRV diagnosis might be misleading at the early stages of the disease [11]. Symptoms of RRD are diverse and include: red abnormal coloration of shoots and foliage, sprout elongation, excessive thorn proliferation, witches broom, which has been also associated with herbicides glyphosate [11], internodes shorten (stem shorter than leaves), dwarfed, elongated or misshapen leaves, flowers may be mottled or distorted, thick secondary branches compared to primary branches, chlorosis on leaves reduced resistance to winter, and death of the infected plants, which may occur in 1 or 2 years [11, 12]. Apparently, rose plants lose resistance to winter conditions.

Early virus detection by symptoms is challenging due to common multiple virus infections in roses, and putative characteristics of the rose tissue, which can be more laborious to process if compared to diagnostic procedures in use for other phytopathogens. The selection of the detection methods applied to plants, or virus vectors, play an important role in the final diagnostic output and management of viral diseases [13, 14]. One of the most utilized techniques is RT-PCR (Reverse Transcription-Polymerase Chain Reaction) which exponentially amplifies target-specific diagnostic sequences. The RT-PCR reaction mix is incubated through cycles of different alternating temperatures (denaturation, annealing, and extension) requiring the assistance of a thermal cycler [15]. Alternatively, isothermal DNA amplification methods amplify the targeted diagnostic sequence at a constant temperature. Allowing to replace thermal cyclers with more simple equipment able to maintain a constant temperature, and output characteristics as PCR reactions such as LoD and specificity [16].

Loop-Mediated Isothermal Amplification (LAMP) is isothermal method that achieves a high degree of specificity and efficiency. LAMP uses a DNA polymerase and a set of four to six oligonucleotides specifically designed to recognize four to six regions of the diagnostic target DNA [16]. In addition, LAMP results can be visualized at the naked eye using either a pre-reaction and pH-sensitive dye (Hydroxy Naphtol Blue) or a post-reaction fluorescent dye (PicoGreen or SYBR Green I) [17, 18].

Among, the best management practices once the RRD occurs is to remove symptomatic plants including the root ball seeking to minimize secondary inoculum and the spreading of viruliferous mites to healthy plants, which in time causes economic losses [1]. Accurate, sensitive, and field transferable detection of RRV is needed for early detection of RRV in foundation blocks, susceptible breeding lines, and biosecurity. This study hypothesizes the technological need can be fulfilled by exploring the development of a LAMP method while investigating chemistries and dyes available for naked eye visualization.

## 2. Material and Methods

A flow chart showing the RRV genes targeted and the sequence of methods and chemistries explored is shown in Fig 1.

**Fig 1.**
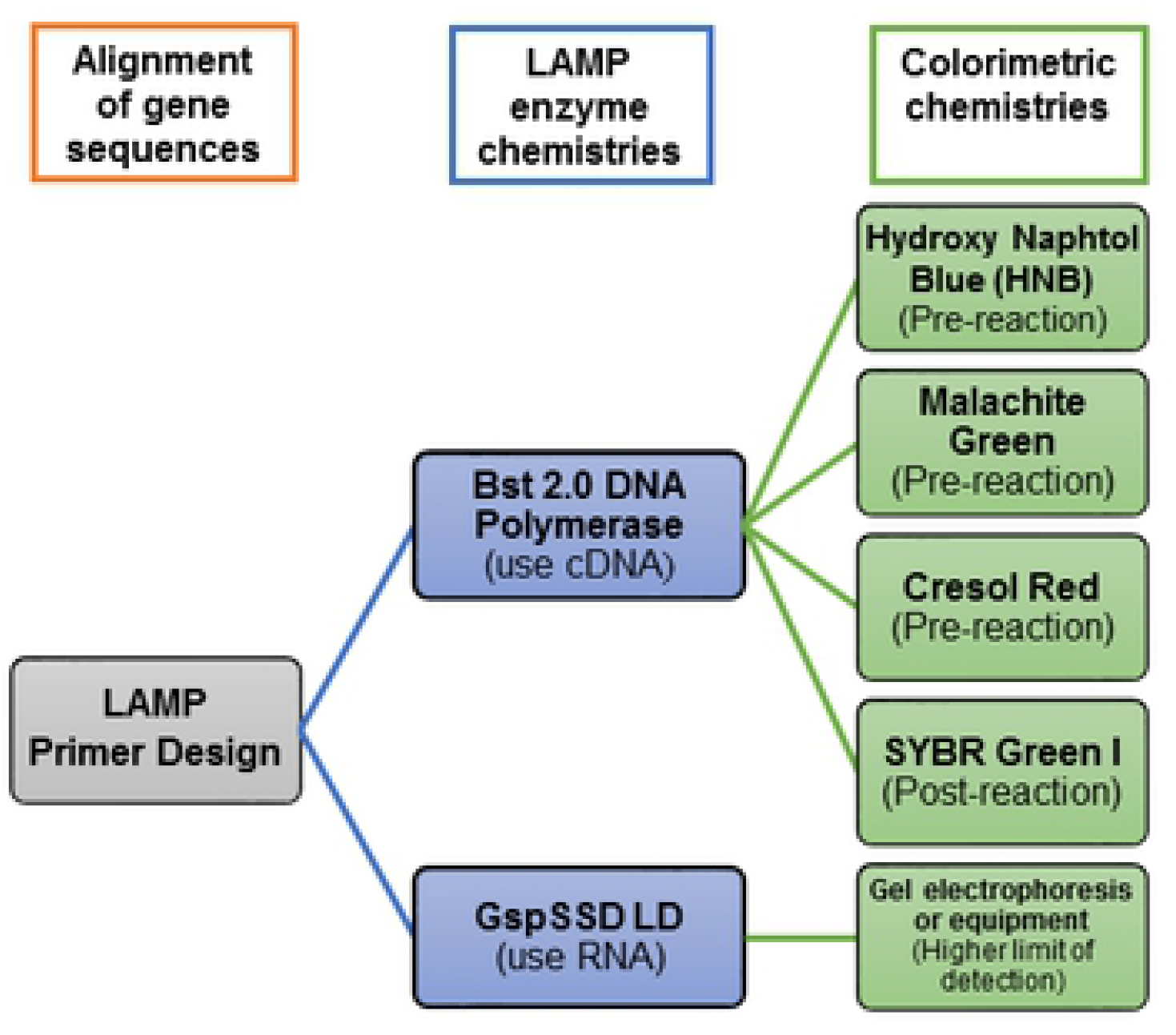
Flow chart showing the steps, methods, and chemistries used to explore Reverse Transcription Loop-Mediated Isothermal Amplification (RT-LAMP) targeting two genes of rose rosette virus for detection of rose rosette disease.

### 2.1. Source of viruses

Infected plant tissue was from Oklahoma where RRV is endemic. Samples were sourced by the Plant and Insect Disease Diagnostic Laboratory (PDIDL), Oklahoma State University (OSU), and were collected from two RRV resistance rose-varietal plots located in Perkins and the Tulsa rose garden, Oklahoma. Nine lyophilized reference-positive control of viruses commonly infecting roses were from Agdia, Inc (Agdia, Elkhart, IN) and used for specificity assays. These virus species are impatiens necrotic spot virus (INSV), High plains wheat mosaic virus (formerly High plain virus) (HPWMoV), arabis mosaic virus (ArMV), maize stripe virus (MSpV), tomato spotted wilt virus (TSWV), apple mosaic virus (ApMV), prunus necrotic ringspot virus (PNRSV), tomato ringspot virus (ToRSV), and tobacco mosaic virus (TMV). Healthy tissue of *Rosa multiflora* was used as a negative control.

### 2.2. LAMP primer design

Twenty-two RRV nucleoprotein gene sequences (RNA3) were retrieved from the NCBI GenBank. The accession numbers of the analyzed sequences are HQ891913.1, HQ891912.1, HQ891911.1, HQ891910.1, HQ891909.1, HQ891908.1, HQ891907.1, HQ891906.1, HQ891905.1, HQ891904.1, HQ891903.1, HQ891902.1, HQ891901.1, HQ891900.1, HQ891899.1, HQ891898.1, HQ891897.1, HQ891896.1, HQ891895.1, HQ891894.1, HQ891893.1, and HQ891892.1.

Twenty sequences of RRV movement protein gene (RNA4) were retrieved from the NCBI GenBank database. The accession numbers of analyzed sequences are HQ891889.1, HQ891888.1, HQ891887.1, HQ891886.1, HQ891885.1, HQ891884.1, HQ891883.1, HQ891882.1, HQ891881.1, HQ891880.1, HQ891879.1, HQ891878.1, HQ891877.1, HQ891876.1, HQ891875.1, HQ891874.1, HQ891873.1, HQ891872.1, HQ891871.1, and HQ891870.1.

The NCBI accessions selected for LAMP primers design were RRV isolates collected in Arkansas, Mississippi, Missouri, Alabama, Tennessee, Iowa, and Oklahoma. Sequences of LAMP oligonucleotides primers were designed using the web interface application Primer Explorer (Eiken Chemical Co., Ltd.) (http://primerexplorer.jp/e/). A set of LAMP primers was selected (Table 1) following parameters described in the Primer Explorer Manual (Table S1). The specificity of the LAMP primers was tested *in silico* using BLASTn [19]. LAMP primers were synthesized by Integrated DNA Technologies (Coralville, IA, USA).

**Table 1.**
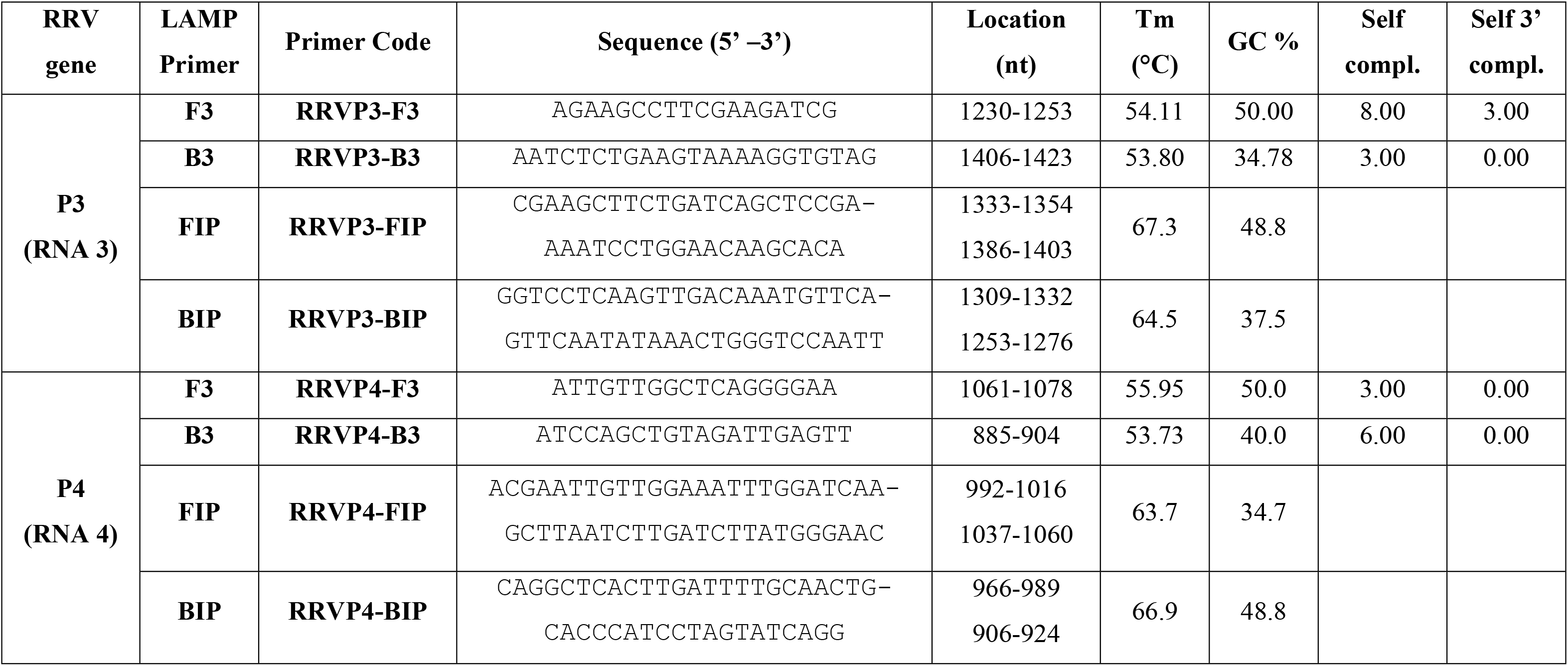
General thermodynamic characteristics of the designed RRV LAMP primers calculated by Primer Explorer.

### 2.3. RNA extraction

RNA extractions were made from either fresh or lyophilized rose tissue and commercially available virus reference controls (Agdia, Elkhart, IN). For fresh tissue, approximately 100 mg of leaves, petals, bark, or roots were loaded in 2 mL microcentrifuge tubes and pulverized using liquid nitrogen and mini-pestels. Total RNA was extracted using the RNeasy plant mini kit (Qiagen Inc., Valencia, CA) following the instructions of the manufacturer. For Agdia lyophilized reference positive controls, 450 µL of RLT buffer (from the Qiagen RNeasy plant mini kit) was directly aliquoted into the positive control vial, vortexed for 30s, and the total RNA extracted following the manufacturer’s instructions.

### 2.4. Virus direct antigen-capture method

Viral RNA was obtained from leaves, petals, bark from young stems, and roots using direct antigen-capture or direct trapping in plastic as described by Babu et al. [20]. Briefly, rose tissue was macerated in 1 mL of phosphate buffer saline (1X PBS) (VWR, Radnor, PA), 0.05% Tween 20, and pH 7.4. Fifty microliters of sap extract were aliquoted into a sterile PCR tube (0.5 ml polypropylene), avoiding air bubbles. The sap was incubated on ice for 2 min, then, the PCR tubes were rinsed twice with 50 µL of 1X PBS-T buffer. To release the viral RNA from the plastic-surface captured virion, 30 µL of nuclease-free water with RNAsin (100 U/mL) (Promega, Madison, WI) was added and incubated at 95°C for 1 min. The supernatant was directly used in reverse transcription reactions.

### 2.5. cDNA synthesis

Four microliters of extracted total RNA obtained either with an RNA extraction kit or the direct antigen-capture method were used. The protocol was performed in two steps, first, denaturation followed by reverse transcription. LAMP RRVP3-B3 or RRVP4-B3 primers (5 µM) were used rather than random hexamer, and Moloney Murine Leukemia Virus Reverse Transcriptase (M-MLV RT) (Promega, Madison, WI), following the manufacturer’s instructions. cDNA synthesis was performed at 37◦C for 90 min.

### 2.6. PCR and cloning of RRV-P3 and RRV-P4 fragments

RT-PCR products were amplified with both LAMP external primers RRVF3-B3 or RRVF4-B4. The PCR reaction mix used was: 10 μL GoTaq Green Master Mix (Promega, Madison, WI), 1 μL RRVP3-F3 and RRVP3-B3 or RRVP4-F3 and RRVP4-B3 primers (5 μM), 3 μL of cDNA template, 1.6 μL of BSA (bovine serum albumin, 50 mg/mL; Ambion, Austin, TX), 2 μL 10% PVP40 (polyvinylpyrrolidone, Sigma-Aldrich, St. Louis, MO), and 1.4 μL nuclease-free water (Promega, Madison, WI). The final volume of the PCR reaction was 20 μL. The RT-PCR reaction was performed in a thermal cycler (Biometra, Goettingen, Germany) and cycling parameters were as follows: initial denaturation of 94°C for 5 min, 40 cycles of denaturation at 94°C for 20 s, annealing at 56°C for 20 s, extension at 72°C for 20 s, and a final extension at 72°C for 5 min. The amplified products were analyzed by electrophoresis, 2% agarose gel, in 1X TAE buffer (Tris-acetate-EDTA) stained with SYBR Safe (Invitrogen, Waltham, MA) according to the manufacturer.

The RRV-P3 and RRV-P4 RT-PCR products were excised and purified from the agarose gel, in two stages. First, by QuantumPrep Freeze’N Squeeze Spin Columns (Bio-Rad, Hercules, CA), followed by a second purification with High Pure PCR Product Purification Kit (Roche, Germany). The TOPO TA cloning kit (Invitrogen, Waltham, MA) was used for cloning the purified PCR fragments segments of RRV-P3 and RRV-P4 genes. The TOPO TA kit was used according to the manufacturer’s instructions. Briefly, the purified PCR products were ligated into the pCŔ4-TOPO plasmid and transformed into *Escherichia coli* cells (TOP10, Mach1™-T1R, DH5α™-T1R cells) following the manufacturer’s protocol. Plasmids were stored at -20°C until use.

### 2.7. LAMP optimization

The purified plasmids carrying the RT-PCR inserts of RRV-P3 and RRV-P4 were used for optimization purposes. The optimal temperature was assessed from a gradient of temperature assay from 60°C to 72°C and the optimal MgSO4 concentration was determined by testing nine different concentrations from 2mM to 10mM.

LAMP reagents were: 2.5 μL isothermal amplification buffer (1X; Biolabs, Ipswich, MA), 2.5 μL betaine (0.8 M; Lucigen, Middleton, WI), 1 μL MgSO4 (4 mM; Biolabs, Ipswich, MA), 1.4 μL dNTPs (1.4 mM, GeneScript, Piscataway, NJ), 1 μL LAMP primers F3 – B3 (0.2 μM), 1 μL AMP primers FIB – BIP (0.8 μM), 1 μL Bst 2.0 WarmStart^®^ DNA polymerase (0.32 u; Biolabs, Ipswich, MA), 1 μL Hydroxy Naphthol Blue (HNB) (120 μM; Sigma-Aldrich, St. Louis, MO), 1 μL RRV-P3 plasmid or 3 μL of cDNA from rose tissue or nuclease-free water (Promega, Madison, WI). The final volume of the LAMP reaction was 25 μL.

LAMP was performed in a dry bath incubator (GeneMate/Bioexpress, Kaysville, UT). LAMP reactions for RRV-P3 were incubated for 1 h at 64°C and 1h at 66.5°C for RRV-P4. The polymerase deactivation was performed at 80°C for 10 min, at the end of the reaction. The amplified products were analyzed by electrophoresis in 2% agarose gel, 1X TAE buffer, and stained with SYBR Safe (Invitrogen, Waltham, MA). The colorimetric reaction was optimized by adding 1 µL HNB (120 µM, pre-reaction) or 3 µL of freshly prepared 10-fold dilution of SYBR Green I dye post-reaction(Invitrogen, Waltham, MA) after amplification of each LAMP reaction (Fig 1). Cresol red and Malachite green (Sigma-Aldrich, St Louis, MO) were also tested to assess their colorimetric reaction using Bst 2.0 WarmStart^®^ DNA polymerase.

### 2.8. Quantitative LAMP

Quantitative LAMP (qLAMP) reactions were carried out in 15 µL isothermal GspSSD2.0 master mix (ISO-004) (OptiGene, Horsham, UK), 1 μL LAMP primers F3 – B3 (0.2 μM), 1 μL LAMP primers FIB – BIP (0.8 μM), and 1 μL RRV-P3 plasmid or 3 μL of RNA extracted from rose tissue. Nuclease-free water was used for non-template control (Promega, Madison, WI). The reaction was performed in a Rotor-Gene 600 thermocycler (Corbett Research, Qiagen Inc., Valencia, CA) at 64°C for 50 min. Enzyme deactivation was performed at 80°C for 10 min at the end of the reaction. The reaction data were registered and analyzed using the Rotor-Gene 6000 series and software 1.7 (Corbett Research, Qiagen Inc., Valencia, CA).

### 2.9. Limit of detection and specificity assays

LoD assays were performed using ten-fold serial dilutions from 1 ng/µL to 1 fg/µL of plasmids RRV-P3 and RRV-P4. The plasmid concentration was quantified using a Nanodrop 2000 (Thermo Scientific, Waltham, MA). One microliter of each dilution was used as a template for LAMP.

For specificity, the LAMP primers were tested with RNA extracted from nine lyophilized reference virus controls (Agdia, Elkhart, IN) listed above in the section ‘source of viruses’. Plasmid RRV-P3 (positive control), cDNA and RNA from healthy rose tissue (negative control), and nuclease-free water (non-template control) were included in both LoDs and specificity assays. The results of these assays were analyzed in 2% agarose electrophoresis in 1X TAE buffer, colorimetric reaction, and qLAMP.

### 2.10. Screening of field samples

RRV-P4 LAMP was tested with 38 field samples including symptomatic and non-symptomatic rose samples collected at the Oklahoma State University RRV resistance rose varietal plot by PDIDL. RRV-P3 LAMP was tested with 30 samples of rose tissue collected at the Tulsa rose garden. The LAMP reaction conditions were according to optimized parameters tested and described above.

Endpoint RT-PCR was performed as a reference and confirmatory assay using the RRV primers and cycling conditions described by Dobhal et al. [11]. Briefly, 10 μL GoTaq Green Master Mix (Promega, Madison, WI, USA), 1 μL RRV2F (5’-TGCTATAAGTCTCATTGGAAGAGAAA-3’) and RRV2R (5’-CCTATAGCTTCATCATTCCTCTTTG-3’) (5 μM) primers, 3 μL of cDNA of rose samples, 1.6 μL BSA (50 mg/ml; Ambion, Austin, TX), 2 μL PVP 40 (10%; Sigma-Aldrich, St. Louis, MO), and 1.4 μL nuclease-free water (Promega, Madison, WI). The final volume of the reaction was 20 μL.

RT-PCR was performed in a thermal cycler (Biometra, Goettingen, Germany), the amplified products were visualized in 2% agarose gel electrophoresis in 1X TAE buffer and stained with SYBR Safe (Invitrogen, Waltham, MA) according to the instructions of the manufacturer. The PCR cycling consisted of initial denaturation of 94°C for 3 min followed by 38 cycles of denaturation at 94°C for 20 s, annealing at 56°C for 30 s, extension at 72°C for 30 s, and final extension at 72°C for 3 min.

## 3. Results

### 3.1. LAMP primer design

The primers were selected after analysis of the thermodynamic parameters described by the Primer Explorer manufacturer for selection of LAMP primers (Table S1). LAMP primer sets for RRV-P3 and RRV-P4 aligned with the consensus regions of twenty-two RRV-P3 sequences and twenty RRV-P4 sequences (Table 1). The alignment analysis of outer LAMP primers RRVP3-F3, RRVP3-B3, RRVP4-F3, and RRVP4-B3 was performed *in silico* with BLASTn. The result showed 100% identity with 100% query coverage for both groups of gene sequences of RRV (P3 and P4) analyzed during primer design. Matches with other emaraviruses and virus species were not detected.

### 3.2. LAMP optimization

The sequencing output of the two cloned plasmids carrying the RRV-P3 and RRV-P4 fragments had 99% identity to the RNA3 and RNA4 of RRV isolates in the NCBI database. These fragments were amplified *in vitro* by endpoint RT-PCR with outer LAMP primers RRVP3-F3, RRVP3-B3, RRVP4-F3, and RRVP4-B3. Cresol red and Malachite green did not react in the colorimetric reaction using Bst 2.0 WarmStart DNA polymerase and no change of color was observed.

#### 3.2.1. Primers concentration

Three concentrations of outer LAMP primers RRV-FIP and RRV-BIP at 0.8, 1.6, and 2 μM were tested combined with internal primers F3/B3 and FIP/BIP at ratios 1:4, 1:8, and 1:10 respectively. The best concentration of outer and inner primers 0.2 and 0.8 μM generated the best visible and distinct banding pattern, which is indicative of a positive LAMP reaction amplification (Fig 2).

**Fig 2.**
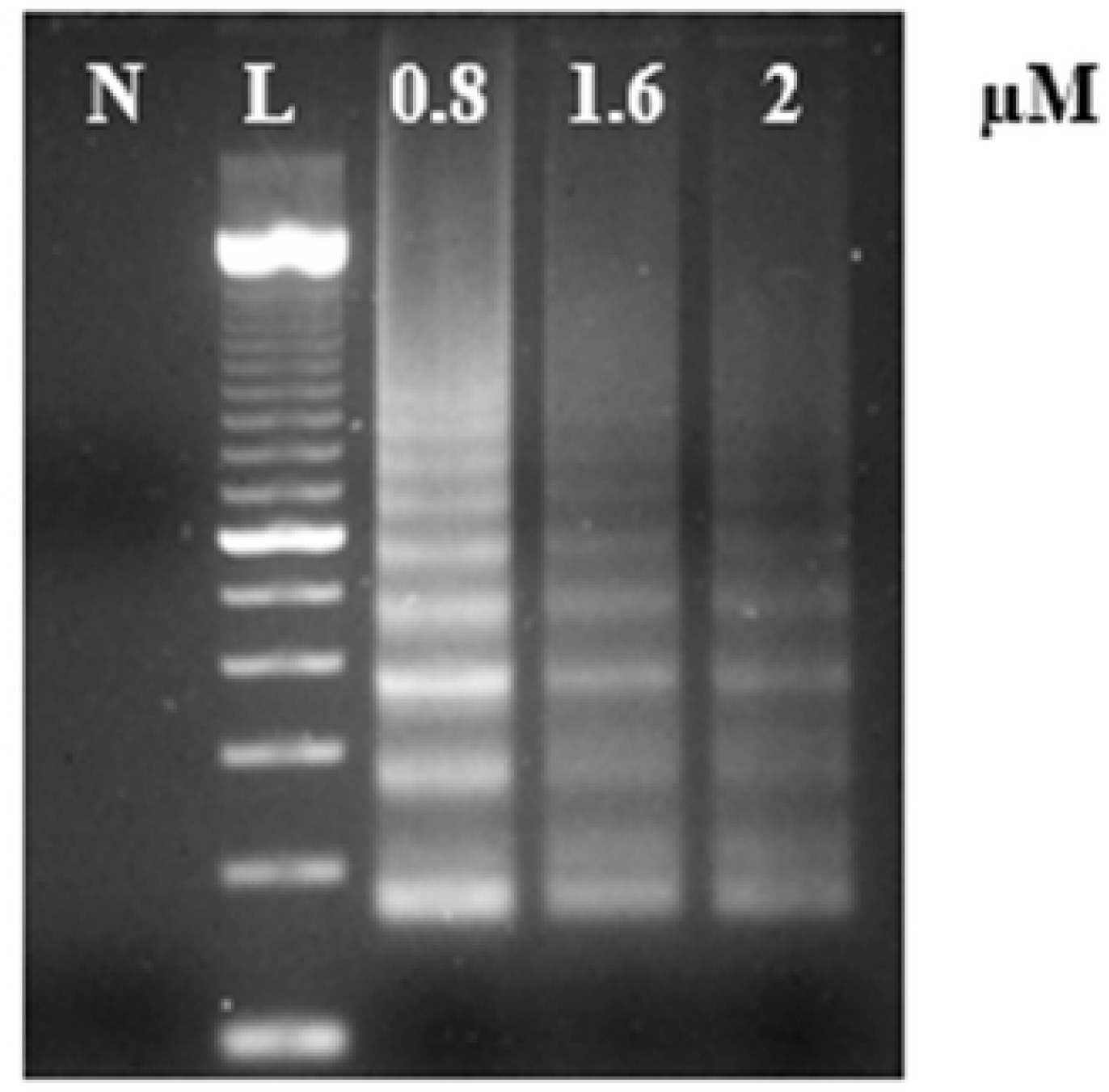
Concentration assay using the outer LAMP primers RRVP3-FIP and RRVP3-BIP, the RRV-P3 plasmid as template, and Bst 2.0 DNA Polymerase. Lane N, is a non-template control (NTC, water); lane L, is a 100 bp DNA ladder.

#### 3.2.2. LAMP performance in temperature gradient

Primer set RRV-P3 amplified the expected diagnostic product within 63 to 68°C (Fig 3A). Similarly, LAMP performed well in a broad range of temperatures amplifying the expected RRV-P4 gene diagnostic product from 60 to 66°C (Fig 3B). The annealing temperature ranged five and six degrees Celsius respectively. Having LAMP performing in a broad range of reaction temperatures reduces errors due to temperature variability that might occur associated with equipment calibration. The amplification temperatures selected for LAMP reactions are 64 ± 1°C for RRV-P3 and 66.5 ± 1°C for RRV-P4 and. No product amplification was detected with the non-template control using the two LAMP primer sets.

**Fig 3.**
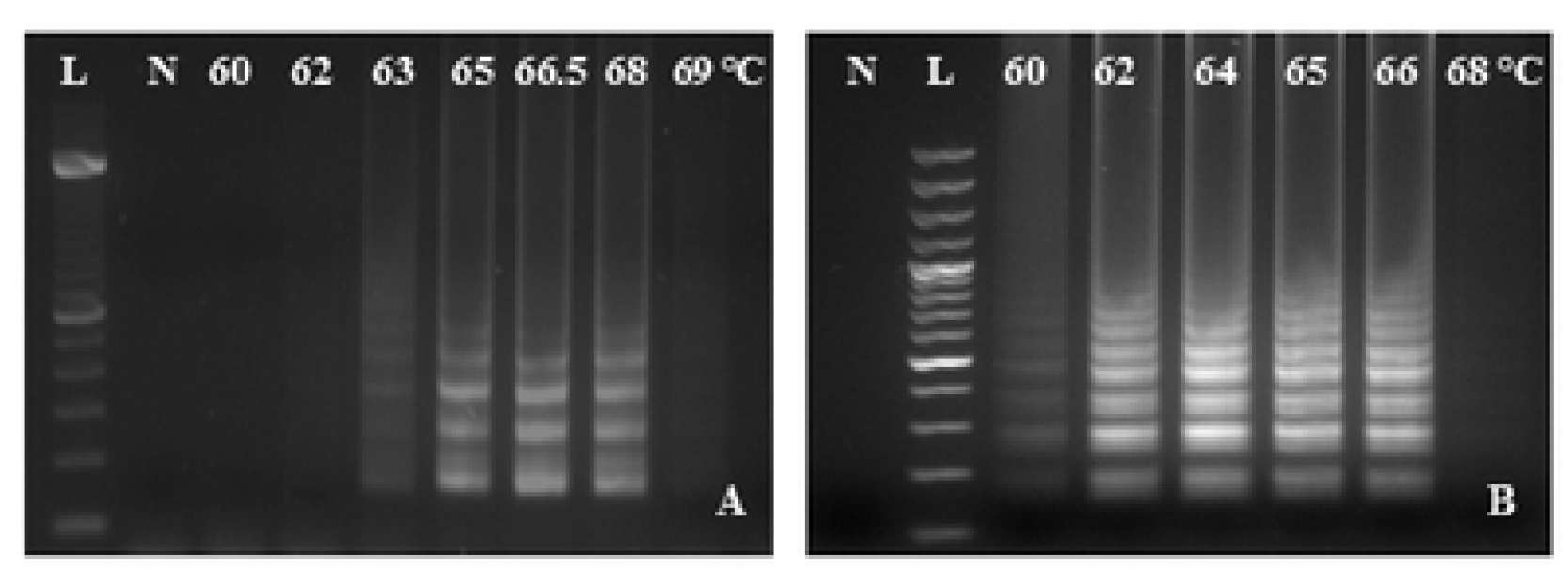
LAMP temperature gradient assay from 60 to 69°C using Bst 2.0 DNA Polymerase. A) RRV-P3 primers and RRV-P3 plasmid as template B) RRV-P4 primers and RRV-P4 plasmid as template. Lane L is a 100 bp DNA ladder and lane N, is a non-template control (NTC, water).

#### 3.2.3. Concentration of magnesium sulfate

The RRV-P3 and RRV-P4 diagnostic products were amplified within six MgSO4 concentrations: 2, 3, 4, 5, 6, and 7mM (Fig 4A) usingBst 2.0 DNA polymerase and the pre-reaction and pH-sensitive dye HNB. The 4mM MgSO4 concentration allowed the amplification and visual discrimination of both RRVP3 and RRVP4 targets. Negative reactions were purple and positive reactions were light blue (Fig 4B).

**Fig 4.**
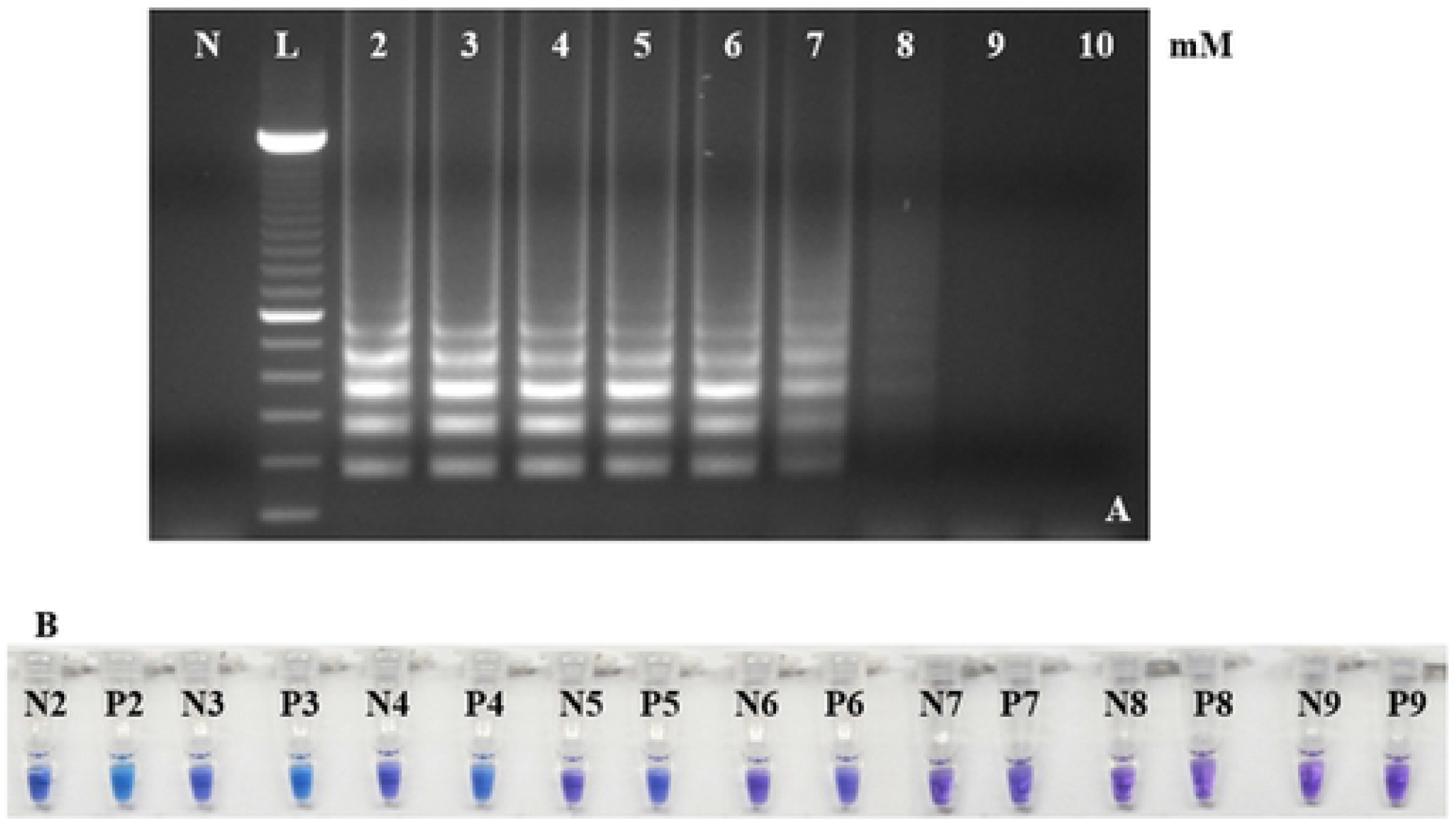
Effect of different concentrations of MgSO_4_ in RRV-P3 LAMP using Bst 2.0 DNA Polymerase. (A) Gel electrophoresis of RRV-P3 LAMP products. Lane 2 is 2 mM, 3 is mM, 4 is mM, 5 is 5mM, 6 is 6mM, 7 is 7 mM, 8 is 8 mM and 10 is 10 mM. Lane N, is a non-template control (NTC, water); lane L, is a 100 bp DNA ladder. The effect of MgSO_4_ concentrations in RRV-P4 LAMP was equal. (B) P tubes are RRV-P3 plasmid (1 ng/µL). P2 to P9 are colorimetric HNB reactions. Tube P2 is 2 mM, P3 is 3mM, P4 is 4 mM, P5 is 5 mM, P6 is 6 mM, P7 is 7 mM, P8 is 8 mM, and P9 is 9 mM. N tubes are non-template controls (NTC, water). Tube N2 is 2 mM, N3 is 3mM, N4 is 4 mM, N5 is 5 mM, N6 is 6 mM, N7 is 7 mM, N8 is 8 mM, and N9 is 9 mM.

### 3.3. LAMP limit of detection

LAMP LoD was tested by serial dilution of cloned RRV-P3 and RRV-P4 amplicons. RRV-P3 and RRV-P4 plasmid concentrations were from 1 ng/µL to 1 fg/µL. The LoD of RRV-P3 using Bst 2.0 DNA Polymerase was 0.1 pg/μL after gel electrophoresis (Fig 5A). The colorimetric LoD using HNB was 0.01 ng/µL (Fig 5B) and 0.1 pg/ µL with SYBR green I (Fig 5C). The LoD of qLAMP using GspSSD2.0 DNA polymerase was 1 fg/µL (Fig 6). qLAMP allowed quantification of RRV concentration in total RNA extracted from rose tissue, which was 189 fg/µL.

**Fig 5.**
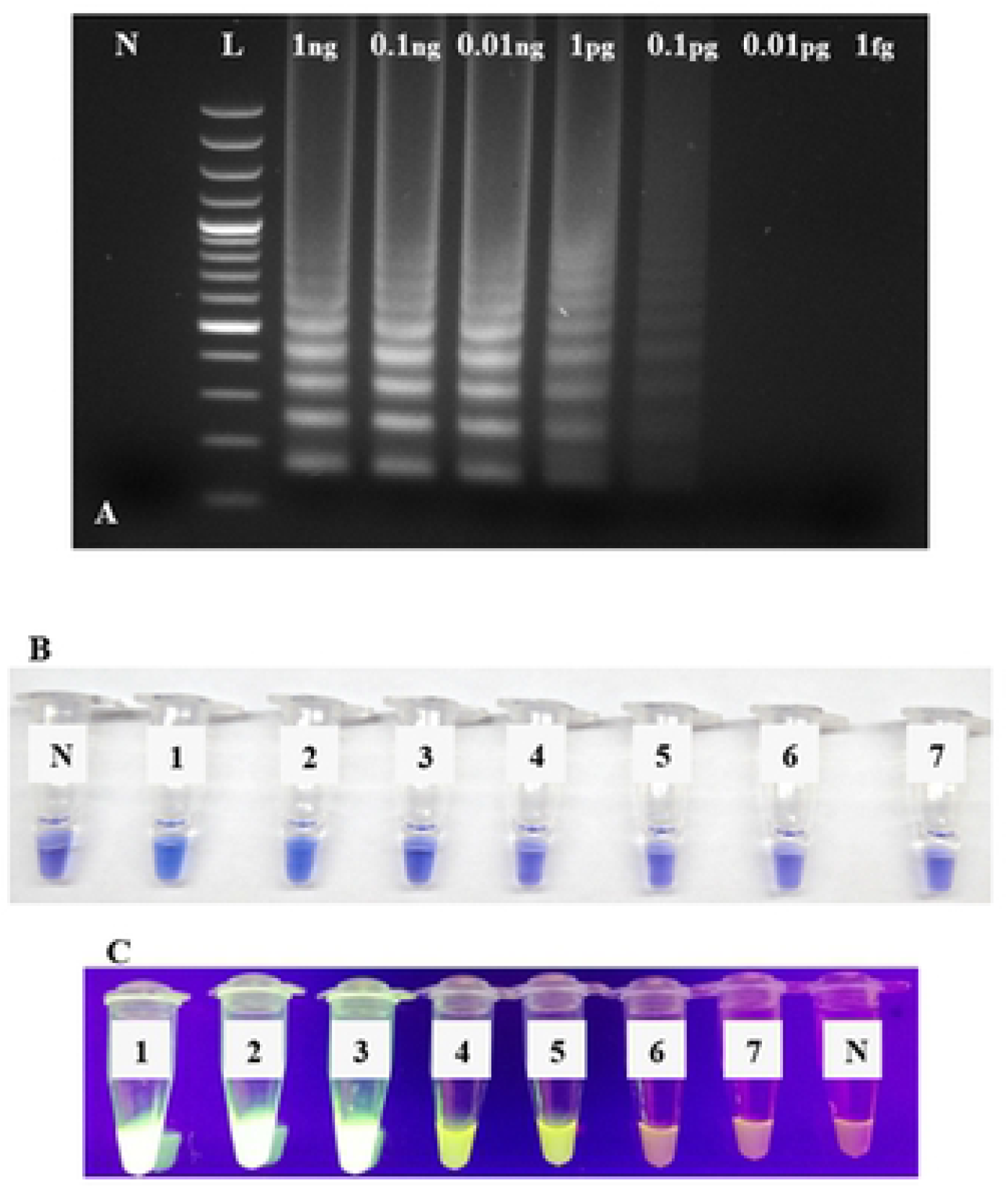
RRV-P3 LAMP LoD assays using Bst 2.0 DNA Polymerase. (A) LAMP LoD by gel electrophoresis using ten-fold serially diluted RRVP-3 plasmid starting from 1ng/µL to 1fg/µL. Lane N, is a non-template control (NTC, water); lane L, is a 100 bp DNA ladder. (B) LAMP colorimetric LoD using with HNB and ten-fold serially diluted RRV-P3 plasmid starting from 1ng to 1fg. Tube N, is a non-template control (NTC, water); tube 1, 1ng/µL; tube 2, 0.1ng/µL; tube 3, 0.01ng/µL; tube 4, 1pg/µL; tube 5, 0.1pg/µL; tube 6, 0.01pg/µL; tube 7, 1fg/µL. (C) LAMP colorimetric LoD using SYBR Green I and ten-fold serially diluted RRV-P3 plasmid starting from 1ng to 1fg. Tube 1, 1ng/µL; tube 2, 0.1ng/µL; tube 3, 0.01ng/µL; tube 4, 1pg/µL; tube 5, 0.1pg/µL; tube 6, 0.01pg/µL; tube 7, 1fg/µL. Tube N, is a non-template control (NTC, water).

**Fig 6.**
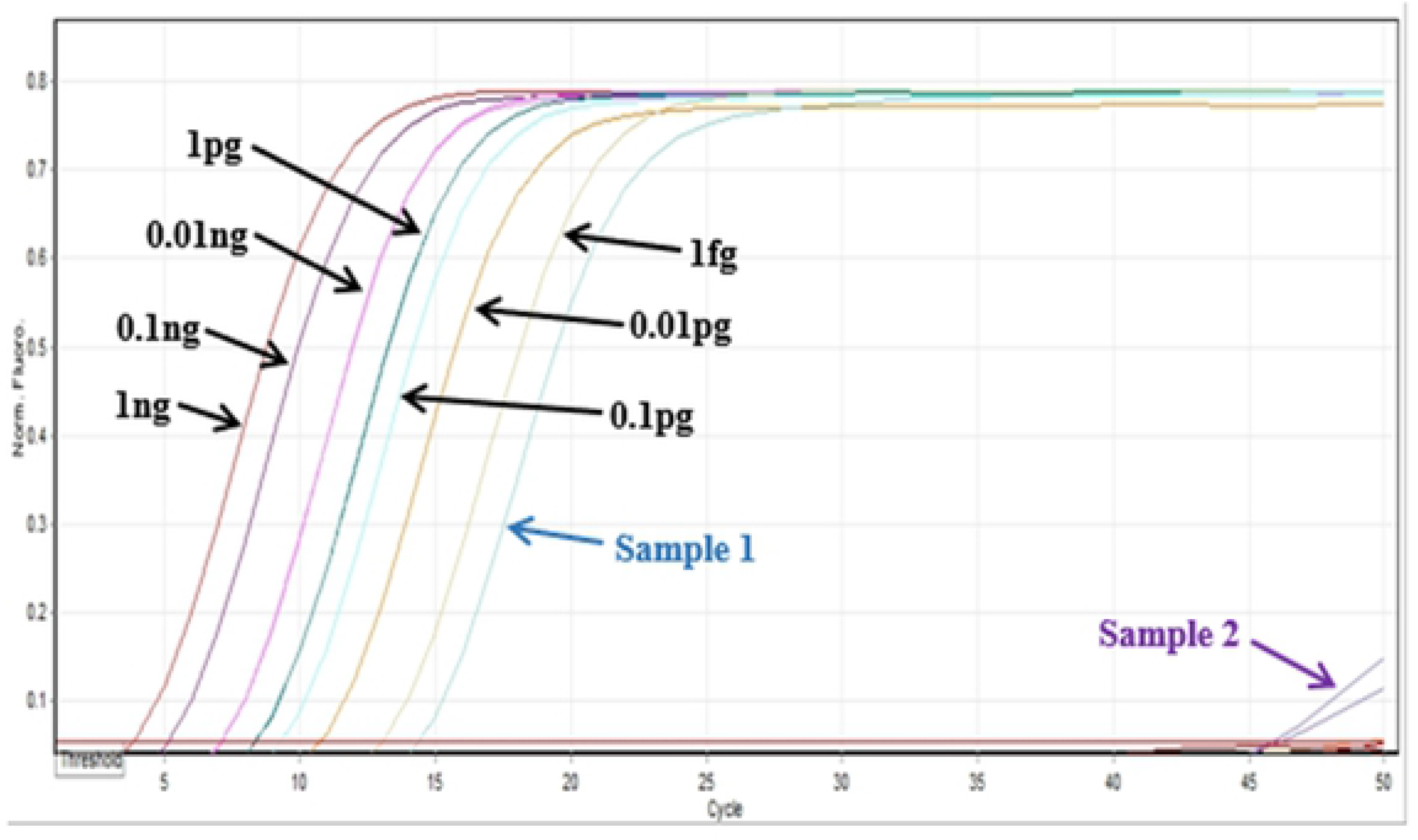
LoD assays q LAMP using GspSSD2.0 DNA polymerase. Limit of detection was made with a ten-fold serially diluted RRV-P3 plasmid starting from 1ng to 1fg and quantification of two unknown RNA from rose tissue were tested (Sample 1 tested positive).

The LoD of RRV-P4 using Bst 2.0 DNA Polymerase is 1 pg/μL (Fig 7A) after gel electrophoresis. The colorimetric LoD using HNB was 0.01ng/µL (Fig 7B).

**Fig 7.**
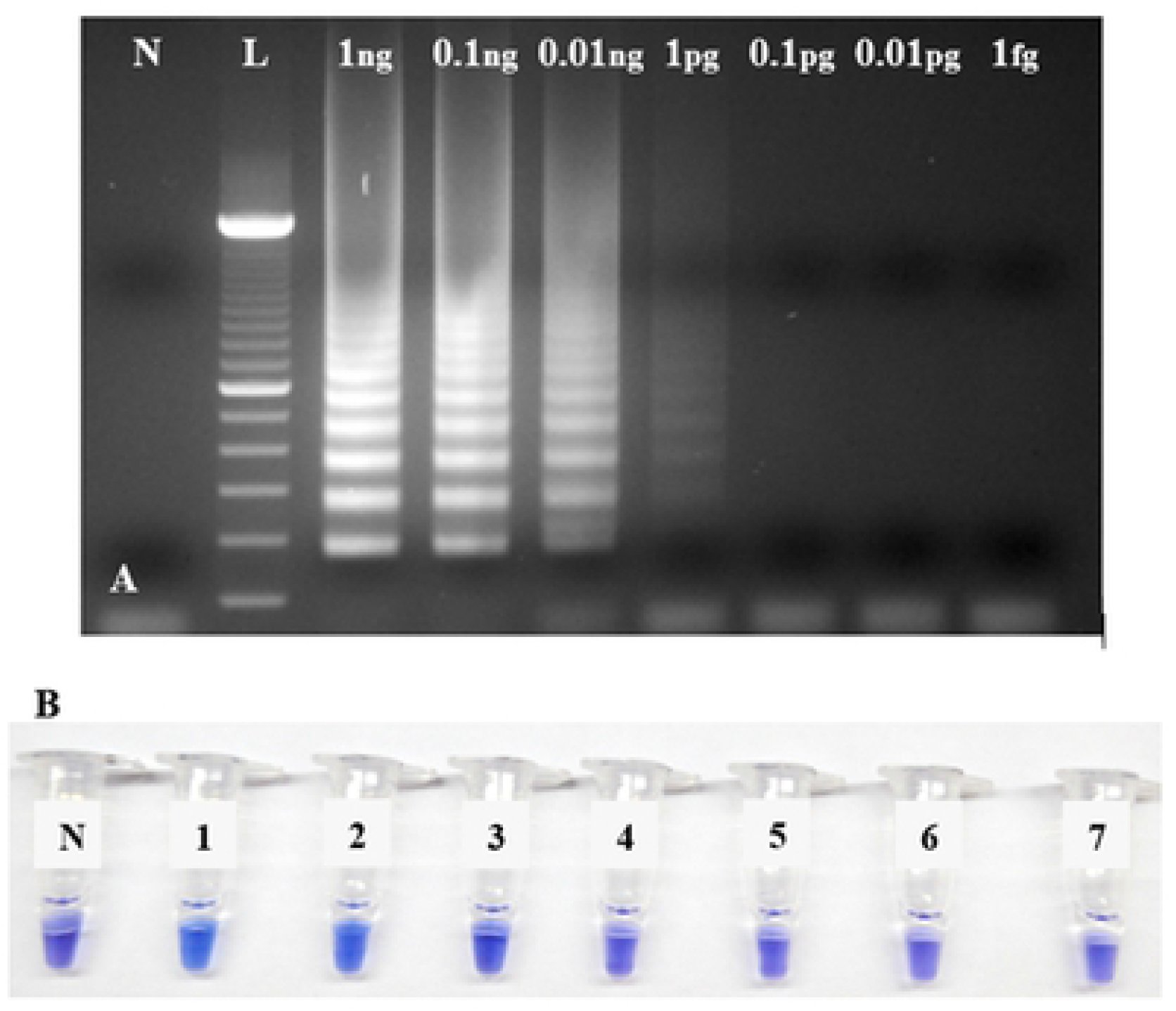
LoD assay of RRV-P4 LAMP performed with Bst 2.0 DNA Polymerase. (A) Gel electrophoresis of RRV-P4 LAMP products from RRV-P4 plasmid serially diluted from 1ng to 1fg limit. Lane N, is a non-template control (NTC, water); lane L, is a 100 bp DNA ladder. (B) Colorimetric RRV-P4 LAMP using HNB. RRV-P4 plasmid was serially diluted from 1ng to 1fg. Tube N, is a non-template control (NTC, water); tube 1, 1ng/µL; tube 2, 0.1ng/µL; tube 3, 0.01ng/µL; tube 4, 1pg/µL; tube 5, 0.1pg/µL; tube 6, 0.01pg/µL; tube 7, 1fg/µL.

### 3.4. LAMP specificity

The specificity of the LAMP primers was tested using the Bst 2.0 DNA Polymerase and gel electrophoresis, HNB, and SYBR green I. No cross-amplification was detected with reference control viruses INSV, HPWMoV, ArMV, MSpV, TSWV, ApMV, PNRSV, ToRSV, and TMV (Fig 8A, 8B, 8C). Similarly, no non-specific amplifications were detected with the cDNAs of these nine viruses using qLAMP (Fig 9). The positive controls targets (1ng/µl of RRV-P3 and RRV-P4 plasmids) were amplified in the three LAMP reactions (Fig 8A, 8B, 8C). The negative control cDNA from healthy rose tissue and the no template control (water) tested negative as expected.

**Fig. 8.**
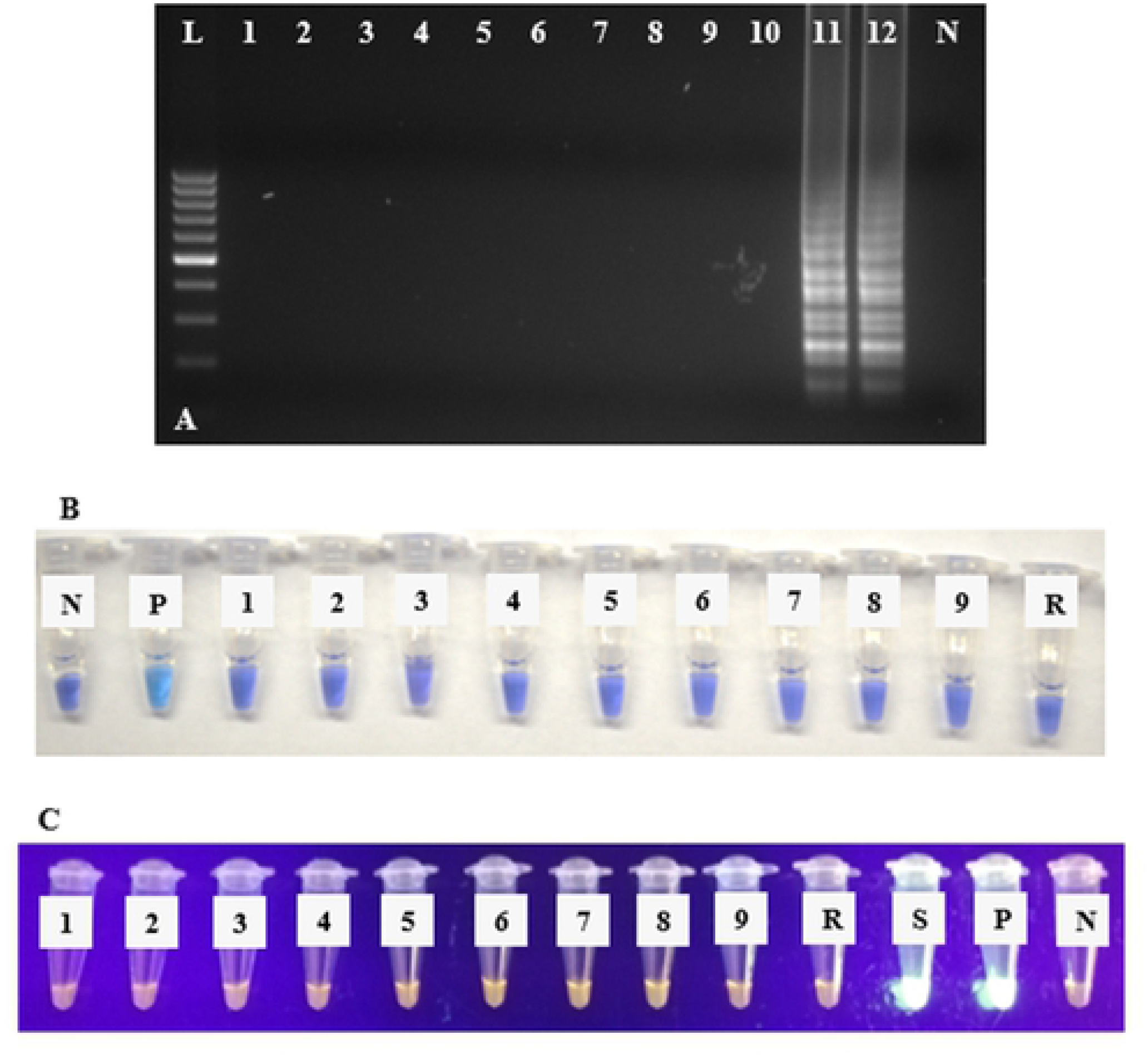
RRV-P3 LAMP specificity assays using Bst 2.0 WarmStart DNA Polymerase using cDNA obtained from nine reference viruses. (A) LAMP gel electrophoresis. Lane L, is a 100 bp DNA ladder, lane 1 is INSV, lane 2 is HPWMoV (formerly High plain virus), lane 3 is ArMV, lane 4 is MSpV, lane 5 is TSWV, lane 6 is ApMV, lane 7, is PNRSV, lane 8 is ToRSV, lane 9 TMV, lane 10 is healthy rose tissue, lane 11 is RRV symptomatic rose tissue, lane 12 is RRV-P3 plasmid, lane N is non-template control (NTC, water). (B) Colorimetric RRV-P3 LAMP **s**pecificity assay using HNB using cDNA of nine reference control viruses. Tube N, is a non-template control (NTC, water), tube P is RRV-P3 plasmid, tube 1 is INSV, tube 2 is HPWMoV (formerly High plain virus), tube 3 is ArMV, tube 4 is MSpV, tube 5 is TSWV, tube 6 is ApMV, tube 7 is PNRSV, tube 8 is ToRSV, tube 9 TMV, tube R, healthy rose tissue. (C) Colorimetric RRV-P3 LAMP **s**pecificity assay using SYBR Green I. tube 1 is INSV, tube 2 is HPWMoV (formerly High plain virus), tube 3 is ArMV, tube 4 is MSpV, tube 5 is TSWV, tube 6 is ApMV, tube 7, is PNRSV, tube 8 is ToRSV, tube 9 TMV, tube R is healthy rose tissue, tube S is RRV symptomatic rose tissue, tube P is RRV-P3 plasmid, and tube N is a non-template control (NTC, water).

**Fig. 9.**
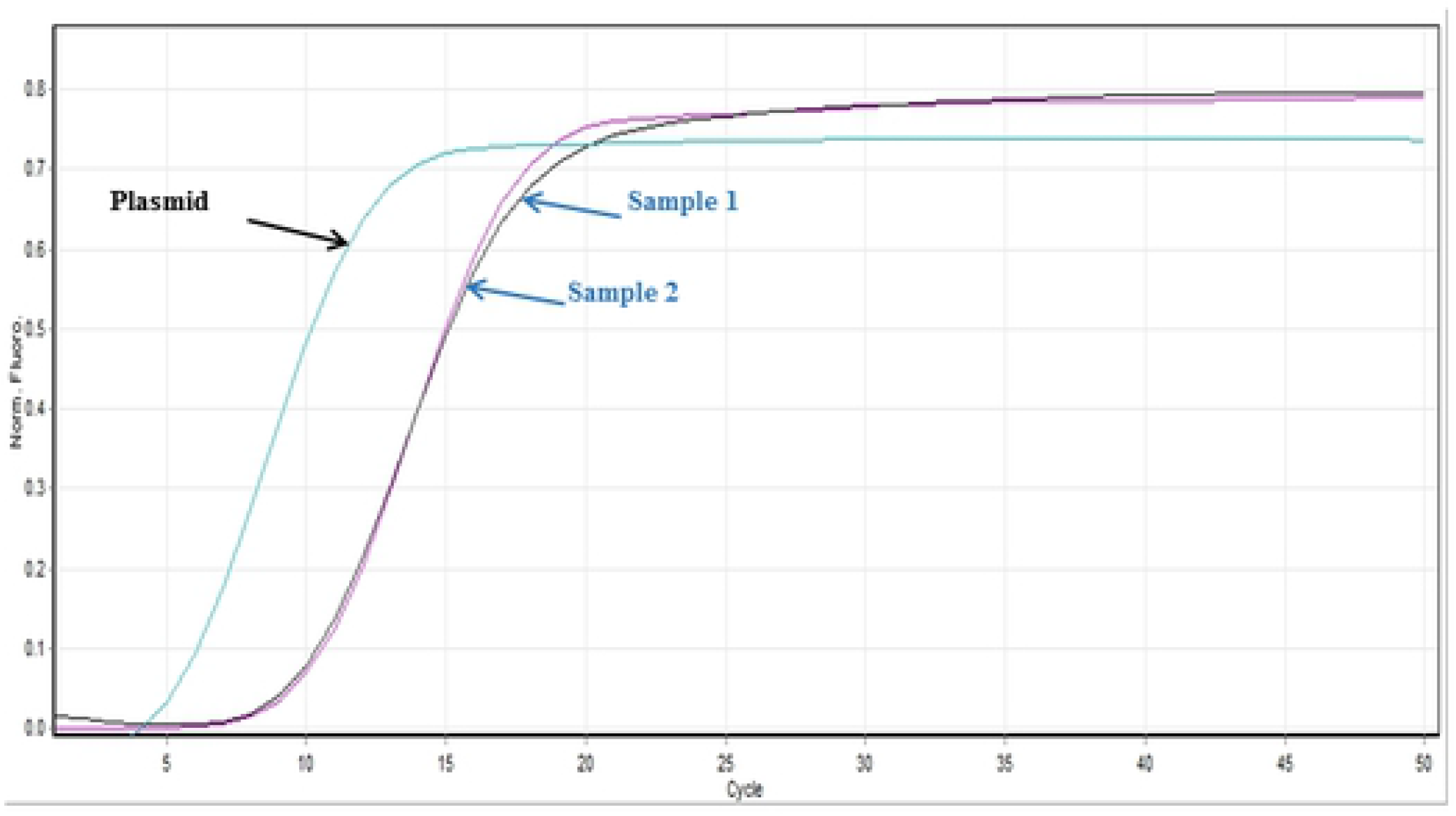
LAMP specificity assays with GspSSD2.0 DNA polymerase. (A) Real-time LAMP specificity using RNA of nine reference control viruses, healthy rose tissue, RRV symptomatic rose tissue (Sample1, 2), and RRV-P3 plasmid.

### 3.5. Screening of infected field samples

A side-by-side comparison between RRV-P4 LAMP with Bst 2.0 DNA polymerase and RT-PCR was done with 38 samples. Twenty-six out of 39 samples tested positive for RRV-P4 LAMP with Bst 2.0 (Table 2). The complete set of 39 samples tested positive by RT-PCR. The AFs BSA and PVP were added to the LAMP reactions and allowed isothermal amplification of the expected RRV-P4 target, however, the colorimetric HNB failed because of unfavorable pH change (Table 2). RNA from rose tissues was extracted using RNeasy Plant Mini Kit in this experiment. The discrepancy between the two methods (13 samples) is due to their differences in LoD, 1pg/µl for LAMP, and 1fg/µl for RT-PCR. The positive control (1ng/µl of RRV-P4 plasmid) tested positive with both RRV-P4 LAMP and RT-PCR while cDNA from healthy rose and the non-template control (NTC) tested negative as expected.

**Table 2.** Side-by-side comparison of RRV-P4 LAMP made with Bst 2.0 DNA Polymerase and RT-PCR. RNA was obtained by virus direct antigen-capture method. Results are from of 38 rose samples from the Oklahoma State University RRV resistance rose varietal plot at Perkins, Oklahoma.

| Nº | Rose Code | Tissue source | LAMP<br>(Bst 2.0 DNA Pol.) | RT-PCR<br>(Dobhal et al., 2016) |
| --- | --- | --- | --- | --- |
| 1 | R1 | Leaves & petals | + | + |
| 2 | R2 | Leaves & petals | + | + |
| 3 | R3 | Leaves & petals | + | + |
| 4 | R4 | Leaves & petals | + | + |
| 5 | R5 | Leaves & petals | + | + |
| 6 | R6 | Leaves & petals | + | + |
| 7 | R7 | Leaves & petals | - | + |
| 8 | R8 | Leaves & petals | - | + |
| 9 | R9 | Leaves & petals | + | + |
| 10 | R10 | Leaves & petals | - | + |
| 11 | R11 | Leaves & petals | - | + |
| 12 | R12 | Leaves & petals | - | + |
| 13 | R13 | Leaves & petals | + | + |
| 14 | R14 | Leaves & petals | - | + |
| 15 | R15 | Leaves & petals | + | + |
| 16 | R16 | Leaves & petals | + | + |
| 17 | R17 | Leaves & petals | - | + |
| 18 | R18 | Leaves & petals | + | + |
| 19 | R19 | Leaves & petals | - | + |
| 20 | R20 | Leaves & petals | - | + |
| 21 | R21 | Leaves & petals | + | + |
| 22 | R22 | Leaves & petals | - | + |
| 23 | R23 | Leaves & petals | + | + |
| 24 | R24 | Leaves & petals | + | + |
| 25 | R25 | Leaves & petals | + | + |
| 26 | R26 | Leaves & petals | + | + |
| 27 | R27 | Leaves & petals | + | + |
| 28 | R28 | Leaves & petals | + | + |
| 29 | R29 | Leaves & petals | + | + |
| 30 | R30 | Leaves & petals | + | + |
| 31 | R31 | Leaves & petals | + | + |
| 32 | R32 | Leaves & petals | + | + |
| 33 | R33 | Leaves & petals | + | + |
| 34 | R34 | Leaves & petals | + | + |
| 35 | R35 | Leaves & petals | - | + |
| 36 | R36 | Leaves & petals | + | + |
| 37 | R37 | Leaves & petals | - | + |
| 38 | R38 | Leaves & petals | - | + |
| 39 | Plasmid RRVP3 | --- | + | Not tested |
| 40 | Healthy rose | Leaves | - | - |
| 41 | NTC | (water) | - | - |

**Note:** (+) samples tested positive for RRV and (-) samples tested negative for RRV.
NTC is non-template control.

The second side-by-side comparison tested 33 samples with RRV-P3 qLAMP using GspSSD2.0 DNA polymerase and RRV-P3 LAMP with BST 2.0 DNA polymerase combined with SYBR Green I for color development post-reaction (Table 3). Twelve samples tested RRV positive using the two methods and five samples tested RRV positive only with RRV-P3 qLAMP using GspSSD2.0. This discrepancy is due to the difference in LoD between qLAMP (1fg/µl) and LAMP using BST 2.0 with SYBR Green I (0.1pg/µl), Fig. 5 and 6. Moreover, 17 samples tested RRV positive using RRV-P3 qLAMP with GspSSD2.0, and thirteen samples tested RRV positive using RRV-P3 LAMP with Bst 2.0 combined with SYBR Green I. Sixteen samples tested RRV negative with these two methods: a) eight samples comprising leaves, bark, and roots were from two varieties (Screaming red neon and Pink surprise). The two varieties were not infected with RRV at the time of sampling. Also, ‘Hybrid 5-13’ samples 2, 4, and 5 (leaves and roots) tested RRV negative with both of the methods but tested RRV positive in the bark. Similarly, ‘Lemon splash’ samples 24 and 26 (stem and roots) tested RRV negative, however, tested RRV positive in the leaves. Another variety, Champlain (samples 19 and 20), tested RRV negative in the leaves but RRV positive in the bark. One out of three specimens of the variety Kiss me rose (samples15-17) tested RRV positive using RRV-P3 qLAMP with GspSSD2.0 in leaves only (sample 15). The obtained results demonstrated the uneven distribution of RRV in rose plants. Positive controls Knock Out’ infected with RRV isolate OK1 and plasmid RRV-P3 (samples 6 and 34 respectively) tested RRV positive with both methods. No amplification or colorimetric reaction was obtained with the negative controls, healthy rose tissue, and non-template control (water) as expected.

**Table 3.** Side-by-side comparison of RRV-P3 real-time LAMP using GspSSD2.0 DNA polymerase and colorimetric RRV-P3 LAMP using Bst 2.0 DNA polymerase with SYBR Green I. Results are from of 33 rose samples from the Tulsa rose garden rose varietal plot, Oklahoma.

| Nº | Rose variety | Tissue source | LAMP (GspSSD2.0) | LAMP (Bst 2.0+SYBR Green I) |
| --- | --- | --- | --- | --- |
| 1 | Caroline Hunt (1) | Leaves | + | - |
| 2 | 5-13 hybrid (1) | Leaves | - | - |
| 3 | 5-13 hybrid (2) | Stem | + | + |
| 4 | 5-13 hybrid (2) | Leaves | - | - |
| 5 | 5-13 hybrid (2) | Roots | - | - |
| 6 | Knock out OK1-IHop (1) | Leaves | + | + |
| 7 | Screaming red neon (1) | Leaves | - | - |
| 8 | Screaming red neon (2) | Roots | - | - |
| 9 | Screaming red neon (2) | Stem | - | - |
| 10 | Screaming red neon (2) | Leaves | - | - |
| 11 | Pink surprise (1) | Leaves | - | - |
| 12 | Pink surprise (2) | Leaves | - | - |
| 13 | Pink surprise (2) | Stem | - | - |
| 14 | Pink surprise (2) | Roots | - | - |
| 15 | Kiss me rose (1) | Leaves | + | - |
| 16 | Kiss me rose (2) | Stem | - | - |
| 17 | Kiss me rose (2) | Roots | - | - |
| 18 | Dulchen (1) | Leaves | + | + |
| 19 | Champlain (1) | Leaves | - | - |
| 20 | Champlain (2) | Stem | + | + |
| 21 | I06-20-14-3x PH (1) | Leaves | + | + |
| 22 | I06-20-14-3x PH (1) | Roots | + | + |
| 23 | Lemon splash (1) | Leaves | + | + |
| 24 | Lemon splash (2) | Stem | - | - |
| 25 | Lemon splash (2) | Leaves | + | + |
| 26 | Lemon splash (2) | Roots | - | - |
| 27 | Above & Beyond (1) | Leaves | + | - |
| 28 | Above & Beyond (2) | Roots | + | + |
| 29 | Above & Beyond (2) | Stem | + | + |
| 30 | Como Park (1) | Leaves | + | - |
| 31 | 5-21 hybrid (1) | Leaves | + | - |
| 32 | Pink double knockout (1) | Leaves | + | + |
| 33 | Apricot drift (1) | Leaves | + | + |
| 34 | Plasmid RRV-3 | --- | + | + |
| 35 | Healthy rose | Leaves | - | - |
| 36 | NTC | (water) | - | - |

**Note:** (+) samples tested positive for RRV, (-) samples tested negative for RRV, (1)
first sampling (2) second sampling. NTC is non-template control.

The third side-by side-comparison contrasts RRV-P3 qLAMP with GspSSD2.0, RRV-P3 LAMP with Bst 2.0 with SYBR Green I, and RT-PCR (Table 4). Varieties Dulchen (sample 7), Pink double knockout (11), Apricot drift (sample12), Lemon splash (sample 14), Above & beyond (samples15,16) tested RRV positive to the three methods. Variety Kiss me Rose and Hybrid 5-21 (samples 6, 10) tested RRV positive only to RRV-P3 qLAMP with GspSSD2.0 and RT-PCR. Varieties Caroline Hunt and Como Park tested RRVpositive to RRV-P3 qLAMP with GspSSD2.0 only. Positive controls plasmid RRV-P3 tested positive with the three methods. No amplification or colorimetric reaction was obtained with the negative controls, healthy rose tissue, and NTC (water) as expected. In general, detection results using RRV-P3 qLAMP with GspSSD2.0 and RT-PCR agreed with the LoD being reported and the varietal results were consistent across tables 3 and 4. The method for direct antigen-capture was consistently trapping RRV directly in plastic from different tissue sources of roses. The addition of SYBR Green I post LAMP reaction generated color change toward fluorescent green in positive amplification, while no change of color or steady-orange was consistent in all negative reactions (no amplification), healthy rose. Amplification and colorimetric reaction (SYBR Green I) was obtained with the RVV-P3 plasmid.

**Table 4.** Side-by-side comparison of RRV-P3 qLAMP using GspSSD2.0, colorimetric RRV-P3 LAMP using Bst 2.0, and RT-PCR. Results are from seventeen field samples of rose from the Tulsa rose garden rose varietal plot, Oklahoma.

| Nº | Rose variety | Tissue source | LAMP (GspSSD2.0) | LAMP (Bst+SYBR Green I) | RT-PCR (Dobhal et al., 2016) |
| --- | --- | --- | --- | --- | --- |
| 1 | Caroline Hunt | Leaves | + | - | - |
| 2 | 5-13 hybrid | Leaves | - | - | - |
| 4 | Screaming neon red | Leaves | - | - | - |
| 5 | Pink surprise | Leaves | - | - | - |
| 6 | Kiss me rose | Leaves | + | - | + |
| 7 | Dulchen | Leaves | + | + | + |
| 8 | Champlain | Leaves | - | - | - |
| 9 | Como park | Leaves | + | - | - |
| 10 | 5-21 hybrid | Leaves | + | - | + |
| 11 | Pink double knockout | Leaves | + | + | + |
| 12 | Apricot dnift | Leaves | + | + | + |
| 13 | Lemon splash | Stem | - | - |  |
| 14 | Lemon splash | Leaves | + | + | + |
| 15 | Above & beyond | Roots | + | + | + |
| 16 | Above & beyond | Stem | + | + | + |
| 17 | Screaming red neon | Leaves | - | - | - |
| 18 | Healthy rose | Leaves | - | - | - |
| 19 | Plasmid RRV-P3 |  | + | + | + |
| 20 | NTC | (water) | - | - | - |

NTC is non-template control.

## 4. Discussion

This study explores, describes, and demonstrates the development of specific, sensitive, and relatively easy-to-use RRV LAMPs. The study used two sets of primers for LAMP amplification of RRV-P3 and RRV-P4 genes which were tested with Bst 2.0 and GspSSD2.0 DNA polymerases. These methods have the potential to be field deployable. LAMP was also tested using SYBR green I and HNB to generate qualitative colorimetric reactions, and a qLAMP was also tested.

The viral nucleic acids, in this case RNA, required for LAMP reactions is commonly extracted from plant tissue using commercially available kits. However, RNA isolated from roses may carry phenolic compounds, starch, other carbohydrates, pigments, and other plant residues in high levels. These compounds inhibit the targeted cDNA and/or DNA amplification because they interact with the viral nucleic acids and reaction mix proteins and cause oxidation and degradation of the RNA [30]. All of which decreases the quality of the samples and causes inconsistent results and concerns about whether all RRV isolates are detectable. Dobhal et al. [31], enhanced RRV RT-PCR detection by adding DNA AFs such as BSA and PVP minimizing the inhibitory effect of putative rose components. However, it was observed the colorimetric LAMP-HNB reaction did not develop a visually distinguishable (sharp) change of color between positive and negative reactions when BSA and PVP were added to the reaction, yet the obtained amplification is detectable by gel electrophoresis. The development of color in colorimetric LAMP-HNB reactions relies on a change in pH occurring during the reaction, and the incorporation of AFs such as BSA and PVP may be interfering with the acidification of the reaction [32].

The direct antigen-capture method reported by Babu et al. [20] allowed rapid and consistent trapping of RRV virions directly on plastic (inner PCR tube surface). The method washes the rose tissue impurities and improves the quality of RNA used in LAMP reactions and it substantially reduces the sample processing time. The method also reduces viral RNA extraction time for LAMP, RPA, and/or RT-PCR to less than 5 min. In summary, the direct antigen-capture method applied to LAMP allowed the RNA reverse transcription and amplification of targeted DNA with no need for BSA and PVP enhancers. Furthermore, the direct antigen-capture did not interfere with pre-reaction pH-sensitive dye (hydroxy naphthol blue) and post-reaction dye (SYBR Green I) in colorimetric LAMP reactions.

The use of only four primers per gene-kit facilitated lamp optimization, assay design, and application of optimized LAMP parameters i.e. optimal outer and inner primer ratios, reaction temperature, and best concentration of magnesium sulfate, which successfully favored the amplification of the expected RRVP3 and RRVP4 targets at 64°C and 66.5 °C respectively. The selected MgSO4 concentration (4mM) allowed steady amplification of both RRVP3 and RRVP4 targets. The optimal concentration of the inner and outer LAMP primers is 0.8µM and 0.2 µM respectively, equivalent to a 4:1 primer ratio. The described Bst 2.0 driven LAMP amplification can be translated to field or nursery testing if using the fluorescent (SYBR green I). The pH-sensitive dyes (HNB) is less recommended. An advantage of LAMP colorimetric results is that can be judged either at daylight or UV light. A critical difference to consider if pursuing field-testing is that SYBR Green I has to be added at the end of the LAMP reaction, while HNB must be added to the mix before the LAMP reaction starts (Fig 1). SYBR green I allowed clear visual discrimination between infected and healthy samples compared to HNB (Figs 5B, 6B, and 6C). The visual LoD of SYBR green I and HNB are 0.01ng/μL and 0.1pg/Ll, respectively.

The LoD between the two LAMP chemistries studied was different. Bst 2.0-based LAMP detected RRV to 1pg/μL while GspSSD2.0 qLAMP allows detection to1fg/μL. The qLAMP assay from this study and the RRV RT-PCR reported by Dobhal et al. [11] have equivalent LoDs (1fg/μL) (Fig 8).

The two RRV-LAMP sets of primers did not cross-react with eight reference control virus commonly co-infecting rose viruses, one taxonomically related emaravirus to RRV, and healthy rose tissue, which confirmed the predicted specificity pairwise analysis results obtained *in-silico* using Primer-Blast and Blastn. In terms of specificity, no difference between LAMP primer sets RRV-P3 gene (viral coat protein) and RRV-P4 gene (movement protein) was detected (Fig 9).

The qLAMP assay is relatively less time-consuming because was combined with the direct antigen-capture or direct trapping in plastic which takes 10-15 minutes for a batch of 1-15 samples. The GspSSD2.0 qLAMP reaction takes approximately 1 hour and uses RNA directly as template while RT-PCR reaction time takes circa 2 hours and uses cDNA as a template. cDNA requires an additional 45 min to 1 hour. In this research, GspSSD2.0 qLAMP was performed with a thermocycler (RotorGene 6000), but fluorescence recordings can be also performed using a simpler fluorescent reader that allows on-site testing of doubtful symptomatic rose plants. The side-by-side comparison of LAMP and RT-PCR [11] showed inconsistent detection of RRV by RRV-P4 LAMP with Bst 2.0-polymerase if compared to RT-PCR (Table 2). Twelve samples tested negative out of 38 RT-PCR positives. The inconsistencies are explained by the lower LoD reported for RRV RT-PCR [11]. Improvement in LAMP primer design would improve the LoD of the method. A second side-by-side comparison between RRV-P3 qLAMP using GspSSD2.0 and the colorimetric RRV-P3 LAMP using Bst 2.0 combined with SYBR Green was performed with 33 rose samples (Table 3). Twelve samples tested RRV positive using the two methods and five samples tested RRV positive only with RRV-P3 qLAMP GspSSD2.0 because RRV-P3 LAMP Bst 2.0 combined with SYBR Green I has a higher LoD. (qLAMP 1fg/µl and LAMP using BST 2.0 + SYBR Green I 0.1pg/µl). Another group of 17 samples tested positive using RRV-P3 qLAMP with GspSSD2.0, and thirteen samples tested positive using RRV-P3 LAMP with Bst 2.0 combined with SYBR Green I, other 16 samples tested negative with these two methods. Testing leaves, bark, and roots from different varieties demonstrated the uneven distribution of RRV in rose plants which is detailed in Table 3.

A third side-by-side comparison of RRV-P3 qLAMP GspSSD2.0, colorimetric RRV-P3 LAMP Bst 2.0 with SYBR Green I, and RT-PCR was made targeting seventeen rose samples collected at the RRV resistance rose varietal plot located at the Tulsa rose garden (Table 4). also shows discrepancies in detection among the three methods. The GspSSD2.0 detected 2 RRV positives out of the 17 samples that were not detected by the colorimetric Bst 2.0 DNA Polymerase plus SYBR Green I and RT-PCR. These results are consistent with the low LoD of Bst 2.0 being reported and the varietal results were consistent across tables 3 and 4.

DNA observed in this research and the uneven distribution of RRV in the plant which may cause RRV to pass not adverted by RT-PCR. In general, RRV was detected from leaves, petals, stem (bark) and roots. Consistent amplification of RRV was obtained from leaves, however, as noted inconsistencies using RT-PCR point toward the uneven distribution of RRV in their hosts and the need for a statistically based sampling method. In general, the obtained LAMP results agree with developed LAMPs reported for plant viruses such as banana bunchy top virus (BBTV), banana streak viruses (BSVs), cucumber mosaic virus (CMV), tomato chlorosis virus (ToCV), potato virus Y (PVY), tobacco etch virus (TEV), tobacco mosaic virus (TMV), and rice ragged stunt virus (RRSV). These LAMPs used the Bst 2.0 DNA polymerase [21–23]. Another polymerase, GspSSD2.0 was also reported for LAMPs assays for turnip yellows virus (TuYV), chrysanthemum stem necrosis virus (CSNV), fig mosaic virus (FMV), little cherry virus (LChV), and sugarcane mosaic virus (SCMV) [24–29].

## 5. Conclusion

This study explored LAMP with two DNA polymerases, qLAMP with GspSSD2.0 confirmed by gel electrophoresis, and colorimetric LAMP with Bst 2.0. Four different dyes were tested, but visible colorimetric reactions were obtained only when LAMP Bst 2.0 was combined with SYBR I or HNB. The last is less recommended since the change of color is low contrast. qLAMP with GspSSD2.0 offers LoD equal to RT-PCR, takes a shorter time since it works with RNA directly, however, does not support colorimetry. LAMP with Bst 2.0. does not have an LoD as low as RT-PCR, but it supports colorimetry. The colorimetric LAMP Bst2.0 LoD can be improved through an oligonucleotide primer design. Colorimetric LAMP Bst2.0 also takes more reaction time since requires a reverse-transcription step. In general, the isothermal LAMPs tested has potential for field application and monitoring of virus-free germplasm in nurseries, selection of RRV-resistant germplasm, also biosecurity at points of entry, and microbial forensics.

## 6. Acknowledgements

This research was funded by the USDA, NIFA, SCRI grant ‘Combating Rose Rosette Disease: Short Term and Long-Term Approaches’ (2014-51181-392 22644/SCRI), and the Oklahoma Agricultural Experiment Station, Hatch # Okl 02950 ‘Detection and Diagnostic Methods For Agricultural Biosecurity and Forensic Plant Pathology Applications’. We acknowledge Dr. Binoy Babu and Mathews Paret (University of Florida) for sourcing a protocol for viral RNA extraction before publication, Dr. Mohammad Arif (University of Hawaii) for reviewing the manuscript, Optigene for sourcing free samples of GspSSD Mastermix ISO-004, and the Oklahoma State University Biochemistry and Molecular Biology Core Facility for assistance and cooperation. The mention of trade names or commercial products does not imply a recommendation or endorsement by the authors or Oklahoma State University. No conflict of interest exists.

## 8. Supporting information

S1 Table. Primer Explorer selection parameters for LAMP primer set.

